# New Insights into Epileptic Spasm Generation and Treatment from the TTX Animal Model

**DOI:** 10.1101/2025.02.25.639129

**Authors:** John W. Swann, Carlos J. Ballester-Rosado, Chih-Hong Lee

## Abstract

Currently, we have an incomplete understanding of the mechanisms underlying infantile epileptic spasms syndrome (IESS). However, over the past decade, significant efforts have been made to develop IESS animal models to provide much-needed mechanistic information for therapy development. Our laboratory has focused on the TTX model, in which tetrodotoxin (TTX) is infused into the neocortex of infant rats, producing a lesion at the infusion site and mimicking IESS resulting from acquired structural brain abnormalities. Subsequent electrophysiological studies showed that the epileptic spasms originate from neocortical layer V pyramidal cells. Importantly, experimental maneuvers that increase the excitability of these cells produce focal seizures in non-epileptic control animals but never produce them in TTX-infused epileptic rats; instead, epileptic spasms are produced in epileptic rats, indicating a significant transformation in the operations of neocortical networks. At the molecular level, studies showed that the expression of insulin-like growth factor 1 was markedly reduced in the cortex and this corresponded with a loss of presynaptic GABAergic nerve terminals. Very similar observations were made in surgically resected tissue from IESS patients with a history of perinatal strokes. Other experiments in conditional knockout mice indicated that IGF-1 plays a critical role in the maturation of neocortical inhibitory connectivity. This finding led to our hypothesis that the loss of IGF-1 in epileptic animals impairs inhibitory interneuron synaptogenesis and is responsible for spasms. To test this idea, we treated epileptic rats with the IGF-1-derived tripeptide (1-3)IGF-1, which was shown to act through IGF-1’s receptor. (1-3)IGF-1 rescued inhibitory interneuron connectivity, restored IGF-1 levels, and abolished spasms. Thus (1-3)IGF-1 or its analogues are potential novel treatments for IESS following perinatal brain injury. We conclude by discussing our findings in the broader context of the often-debated final common pathway hypothesis for IESS.

We confirm that we have read the Journal’s position on issues involved in ethical publication and affirm that this report is consistent with those guidelines.

**Plain Language Summary:** We review recent findings from the TTX animal model of infantile epileptic spasms syndrome, which show that these seizures come from an area of the brain called the neocortex. In this area, the amount of an important growth factor called IGF-1 is reduced, as is the number of inhibitory synapses that play an important role in preventing seizures. Other results indicate that the loss of IGF-1 prevents the normal development of these inhibitory synapses. Treatment of epileptic animals with (1-3)IGF-1 restored IGF-1 levels and inhibitory synapses and abolished spasms. Thus, (1-3)IGF-1 or an analogue are potential new therapies for epileptic spasms.

---

Epileptic spasms are sudden and brief seizures—only a few seconds in duration—consisting of flexions or extensions of the head, limbs, and/or axial muscles ^1, 2^. They most often occur in clusters and hundreds can occur in a day. This type of epilepsy is most frequently observed during infancy and historically has been referred to as infantile spasms. More recently, the International League Against Epilepsy proposed the term infantile epileptic spasm syndrome, or IESS^3^. However, epileptic spasms can persist beyond infancy in up to 19% of children ^4-6^. Beyond two years of age, these seizures are referred to as epileptic spasms.

Epileptic spasms are unusual seizures whether they occur in infancy or later. Besides being brief, the electroencephalogram (EEG) correlate of the spasms is dominated by an attenuation of the EEG amplitude called an electrodecrement ^7^. A run a of small high frequency spikes can ride the envelope of the electrodecrement. In many instances, the ictal event is initiated by a large generalized slow wave. Between the seizures, a unique chaotic EEG pattern is often recorded in infants, called hypsarrhythmia, which consists of large slow waves that are asynchronous across the cortex and intermixed with large and frequent multifocal interictal spikes ^8, 9^. Treatments used to abolish spasms are different from those for other forms of epilepsy. Widely used anticonvulsants are ineffective but hormone therapies consisting of ACTH or prednisolone are often effective ^2, 10, 11^. Vigabatrin can also be effective, which could implicate GABAergic synaptic inhibition in spasm generation ^12^. Unlike other forms of epilepsy, epileptic spasms and hypsarrhythmia must be eliminated, not merely reduced, for treatment to be judged successful. Using these criteria, 50% of children, at best, will respond to one of these treatments^13^. However, even then, most patients remain neurologically and cognitively impaired, many severely so, especially if treatment is delayed and there is an underlying brain abnormality.

Of all the epilepsies, IESS is considered the most enigmatic ^14^, in part because up to 200 highly diverse clinical conditions are associated with it ^15^. These include etiologies as disparate as perinatal stroke and meningitis, among other insults. In addition, a large and heterogeneous list of single gene mutations has been identified as causative ^16^, yet in a significant percentage of patients, no underlying cause can be identified. How such diverse conditions produce the same unique and stereotyped EEG abnormalities and unusual pharmacological responsiveness has been debated and discussed for decades, without resolution. Nonetheless, it has long been hoped that the development of IESS animal models might elucidate the underlying basic mechanisms, leading to novel disease-modifying therapies. Despite the challenges of reproducing this unique syndrome, several animal models of IESS have been proposed over the past 10-15 years, along with criteria for validating them ^17-23^. Our laboratory has focused its efforts on the tetrodotoxin (TTX) model ^24^.

## The TTX animal model recapitulates clinical epileptic spasms

Like many scientific discoveries, the creation of the TTX model was accidental. In the late 1990s, our laboratory was investigating the role neuronal activity plays in the formation of local cortical networks in early postnatal life. To this end, we sought to block neuronal activity by chronically infusing the sodium channel antagonist tetrodotoxin (TTX) into the developing hippocampus or neocortex ^25^. Unexpectedly, within a week the rats began to display brief seizures with a concomitant EEG discharge. We first considered these events brief focal seizures and focused our efforts on understanding the potential impact of silencing network activity on synapse maturation ^26^. However, a later and more detailed analysis of long-term video EEG recordings showed that the electrographic seizures had the hallmarks of epileptic spasms ^24^. The ictal events of the seizures, which were a few seconds in duration, consisted of a large and generalized initiating slow wave followed by an electrodecrement with fast activity riding its envelope. Spasms often occurred in clusters and were recorded across the neocortex, both ipsilateral and contralateral to the TTX infusion site. The behavioral seizures closely resembled their human counterparts, consisting of brief flexions or extensions of axial and forelimb muscles (for a comparison see: https://www.youtube.com/watch?v=Z4nQxtX3QWk). Importantly, hypsarrhythmia-like or modified hypsarrhythmia-like activity is routinely recorded across the cortex in most epileptic animals, consisting of very large slow waves and frequent and large interictal spikes ^24, 27^. Another criterion proposed for an ideal animal model of infantile spasms is the elimination of spasms after treatment with ACTH and vigabatrin ^28, 29^. In the TTX model, both of these drugs greatly reduced spasm frequency and abolished spasms in 60% of animals treated, mirroring human responsiveness to these treatments. Hypsarrhythmia was similarly eliminated. In addition, TTX-infused animals with spasms have been shown to be learning and memory impaired ^30^.

In the TTX model, spasms are induced at a time comparable to the neonatal period in humans^31^ and behavioral spasms first appear a week later, which is thought to be a time comparable to human infancy ^32^. A potential drawback of the TTX model is that the seizures persist for two to three months and many of our studies have been done on month-old animals. However, as mentioned earlier, while epileptic spasms are most frequently observed during infancy in humans, they can persist beyond infancy in a significant percentage of patients ^5, 6^. Epileptic spasms have also been documented in adolescents and adults with the same electrophysiological and behavioral features as those in infancy ^33-35^. The epileptic spasms in month-old TTX-infused rats parallel these clinical observations. However, there are important differences between the spasms in older rats and older patients: the latter do not respond to ACTH or vigabatrin ^33-35^ and they lack hypsarrhythmia in EEG recordings ^34, 35^. In the TTX model, most animals display hypsarrhythmia-like activity and respond to vigabatrin ^24^ or ACTH ^25^ with a cessation of spasms and hypsarrhythmia. Thus, the TTX model appears to more closely resemble IESS than epileptic spasms in older children and adults. Nonetheless, since so little is known about the mechanisms underlying spasms—whether they occur in infancy or later—the TTX model could provide much-needed information with important implications for infants, older children, and adults with this form of epilepsy.

## Epileptic spasms arise from the neocortex

### Observations in humans

Currently, it is hard to make concrete statements about the area(s) of the brain that generate epileptic spasms in humans, but recent studies have provided new insights. One prevailing view has been that spasms are likely produced by interactions between abnormal cortical and subcortical circuits. The cortex has been proposed as a participant in part because some children with spasms have MRI-diagnosed discrete cortical lesions and no other recognizable brain pathology ^36^. These lesions can arise in various ways. Some result from perinatal brain injuries such as strokes, but others can be from single gene mutations like those that underlie tuberous sclerosis complex. More recently, evidence supporting a cortical origin of spasms has come from epilepsy surgeries. When focal epileptiform EEG activity corroborates a suspected cortical spasm-generating zone, many contemporary pediatric epilepsy surgery programs will now undertake cortical lobectomies, lesionectomies, or hemispherectomies to stop spasms ^37, 38^. A recent meta-analysis of 21 studies reported that nearly 70% of children became seizure-free after cortical resections, strongly suggesting that the observed cortical lesions are responsible for spasms ^39^. In concert with these findings are PET studies reporting focal metabolic abnormalities in the cortex of epileptic spasms patients with normal MRI results^36^. In a portion of these children, spasms have been eliminated when the PET-identified focus was surgically removed ^40-43^.

Nonetheless, other observations have pointed to subcortical contributions to spasms. For instance, another PET study identified hypometabolic regions in the lenticular formation of the globus pallidus ^44^. In a more recent fMRI study of tuberous sclerosis patients with and without IESS ^45^, investigators reported that > 95% of cortical tubers were functionally connected bilaterally to the globus pallidus in infantile spasms patients but not in TSC patients without spasms. Taken together, these findings support the proposition that epileptic spasms likely originate in the cortex and the contributions of subcortical structures such as the globus pallidus are secondary to the cortical discharges^46^.

### Observations in the TTX model

Since video/EEG recordings of the ictal events of epileptic spasms are robust in the TTX model, our laboratory has been able to study their origins. The first indication of a potential cortical contribution to spasms came from the discovery of a cortical lesion at the site of TTX infusion ^29, 47^. This finding was unexpected since TTX is a well-known selective antagonist of sodium channels that generates action potentials. However, previous neurodevelopmental studies had shown that apoptotic programmed cell death in the immature cortex is activity-dependent and that treatments that suppress neuronal activity—like TTX— amplify ongoing neuronal death ^48, 49^. Our subsequent immunohistochemical studies showed a focal region of marked apoptosis at the TTX infusion site in the week after initiating infusion, indicating that apoptosis produced the lesion ^47^. Thus, the TTX model is very likely a perinatal neocortical injury model for spasms.

To pursue potential neocortical mechanisms of spasms, we undertook *in vivo* neurophysiological studies in area S1 of the somatosensory cortex of awake behaving animals^50^. Multielectrode array studies in which current source density and simultaneous multiunit activity analyses were undertaken indicated that the cortex not only actively participated in spasm generation but that the spasms very likely originated from neurons in neocortical layer V. To further explore these findings, we undertook chemogenetic studies to test the hypothesis that the selective activation of layer V pyramidal cells could induce spasms. To this end, we first injected a virus containing the genetic sequence for the DREADDs (designer receptor exclusively activated by designer drugs) hM3D(Gq) into neocortical layer V of epileptic animals and their non-epileptic littermate controls. Since the expression of hM3D(Gq) was under the control of CAM Kinase II, hM3D(Gq) expression was largely restricted to glutamatergic pyramidal cells in layer V ^50^. Next, animals were injected with clozapine to specifically activate DREADDs. Since hM3D(Gq) couples with the G protein-coupled receptor Gq, whose activation is known to result in membrane depolarization and action potential generation, we anticipated that clozapine might induce seizures. Indeed, others had previously shown that the activation of hM3D(Gq) in the neocortex or hippocampus of non-epileptic mice produced behavioral seizures^51^. We obtained similar results in our non-epileptic control rats. As shown in Figure 1, treatment with clozapine resulted in a series of prolonged (10-20 sec duration) subclinical focal seizures at the site of viral transfection. *However, to our surprise, clozapine treatment of epileptic rats never induced focal seizures. Instead, it always produced spasms that were electrographically and behaviorally identical to those that occurred spontaneously during baseline recordings.* We observed a 1.7-fold increase in spasm counts compared to baseline and a 3.5-fold increase in spasm clustering following clozapine ^30^. Thus, chemogenetic results supported the hypothesis that epileptic spasms can be produced by layer V neocortical pyramidal cells and spasms can arise from neocortical networks.

**Figure 1.**
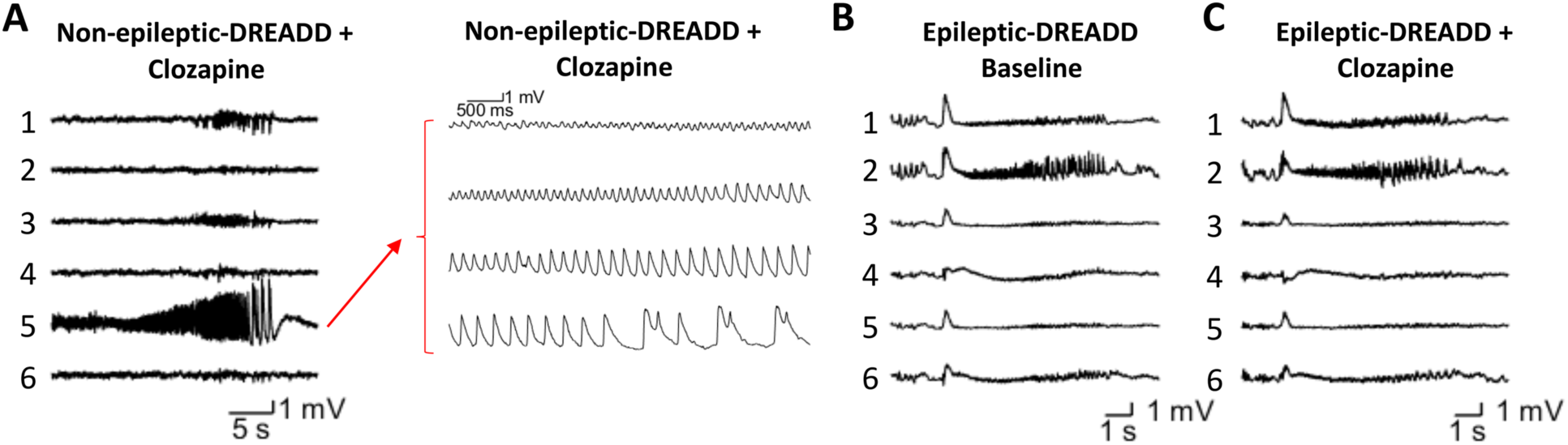
Chemogenetic activation of layer V pyramidal cells induces focal seizures in control animals but spasms in epileptic animals. (A) Left, representative local field potential (LFP) recordings of a focal seizure induced by activating somatosensory pyramidal cells with clozapine at channel 5 in a non-epileptic control animal with DREADD. Right, channel 5 recordings in a faster time base. (B) Recordings of a spontaneous epileptic spasm during baseline recordings in an epileptic animal. (C) Representative LFP traces of an epileptic spasm after clozapine treatment in an epileptic animal. (Modified from reference 50)

Similar results were obtained when we infused the GABAa receptor antagonist picrotoxin into layer V of the somatosensory cortex. Supplemental Figure 1A shows that focal application of picrotoxin in non-epileptic control rats resulted in a prolonged focal seizure. However, using the same experimental protocol in the epileptic rat, picrotoxin never resulted in a focal seizure. Instead, as shown in Supplemental Figure 1C, epileptic spasms were produced that were electrographically and behaviorally identical to those that occurred during baseline recordings (Supplemental Figure 1B). Thus, results were strikingly like those from our chemogenetic studies. Compared to baseline, we observed an eight-fold increase in spasm counts 10 minutes after picrotoxin infusion (Supplemental Figure 1F: 1.57 ± 0.39 versus 0.202 ± 0.05 spasms/10 min). These results strongly support the role of deeper layers of the neocortex in generating epileptic spasms. That said, both studies taken together raise interesting questions. Why didn’t either of the treatments produce focal seizures in animals that have spasms, instead only producing spasms? Does this finding suggest a transformation of the fundamental operations of neocortical networks—that is, that neocortical circuits are rewired in such a way to produce these unique epileptiform and behavioral events?

Finally, while our results clearly implicate the neocortex in epileptic spasm initiation, they do not rule out subcortical contributions. Indeed, we firmly suspect that such contributions are likely since we have observed long delays (230.2 ± 24.3 msec, n=9) between the onset of the initiating slow wave of electrographic spasms and the onset of behavioral spasms in the TTX model. Similarly, clinical investigators have noted hundreds of millisecond-delays between the ictal onset in EEG and electromyographic (EMG) recordings of spasm patients ^52^. Thus, it seems likely that subcortical neuronal networks intercede between the cortex and spinal motor neurons in generating behavioral spasms. Identifying these networks—whether in the globus pallidus, as discussed above, or in brain stem nuclei, as suggested by others ^53-55^—must await future studies.

## Neocortical injury reduces IGF-1 expression and inhibitory interneuron connectivity and implicates them in epileptic spasm generation

Our studies on IGF-1’s role in epileptic spasm generation began in 2012 following a report by Riikonen and colleagues that IGF-1 levels were reduced in the cerebrospinal fluid (CSF) of “symptomatic” IESS patients (i.e., patients with identified etiology and developmental delay at spasms onset) ^56^. By that time, basic science studies of transgenic mouse models had clearly shown that brain-derived IGF-1 played a critical role in normal brain development ^57^. For instance, eliminating IGF-1 or its receptor (IGF-1R) from the brain resulted in reduced brain size, while overexpression of IGF-1 had the opposite effect—enlarged brains ^58, 59^. Previous *in situ* hybridization studies had reported that most, if not all, neurons in the brain express IGF-1 ^60^. Similarly, our findings showed that not only did all neurons in the neocortex express IGF-1, but this growth factor was sequestered in large somatodendritic vesicles ^47^. We were also able to show that IGF-1 is co-expressed with vesicular synaptotagmin 10 ^61^, which others had shown mediated the calcium-dependent release of IGF-1 into the extracellular space ^62^.

When we examined the neocortex of TTX-infused animals with spasms, we observed a marked reduction in the expression of IGF-1, both ipsilateral and contralateral to the infusion site ^47^. One exception to this widespread loss of IGF-1 was in the TTX infusion site itself, where IGF-1 levels were increased. This observation comported with previous studies in other brain injury models showing that reactive astrocytes within a lesion expressed high levels of IGF-1 ^63, 64^. Our biochemical studies further found that signaling through both IGF-1 signaling pathways—IGF-1R-PI3K-AKT and IGF-1R-MAPK—was depressed in epileptic animals ^47^, indicating that the loss of IGF-1 from neocortical neurons impacted downstream signaling through its molecular growth pathways.

While the loss of IGF-1 from the cortex was potentially important in understanding the mechanisms of spasm generation, it alone did not directly implicate the growth factor. Thus, we began to search for a potential mechanism(s) linking IGF-1 to spasms. One clue came from three separate biochemical studies of the CSF of IESS patients ^65-67^. All reported low levels of the inhibitory neurotransmitter GABA and one found the loss only in symptomatic patients ^67^. Other and more recent studies have further implicated GABA in spasm generation. For instance, studies of X-linked infantile spasm syndrome implicated mutations of Aristaless-related homeobox gene (ARX) and a resulting impairment of inhibitory interneuron migration into the neocortex in the genesis of epileptic spasms ^68^. This finding led to the interneuronopathy hypothesis for IESS ^69^. Along these lines, experiments in several animal models of IESS reported deficits in inhibitory interneuron populations in the neocortex ^18, 20, 21, 70-72^. Following these observations, we undertook immunohistochemical studies of glutamic acid decarboxylase, parvalbumin, and synaptotagmin 2 in the TTX model—all biomarkers for inhibitory interneurons and particularly their presynaptic nerve terminals ^61^. Results showed that the expression of all three molecules were markedly depressed in the neocortex of animals with epileptic spasms and coincided with the loss of IGF-1 (Figure 2). And like IGF-1 expression, this loss of inhibitory interneuron connectivity was widespread, occurring both ipsilateral and contralateral to the TTX-induced lesion.

**Figure 2.**
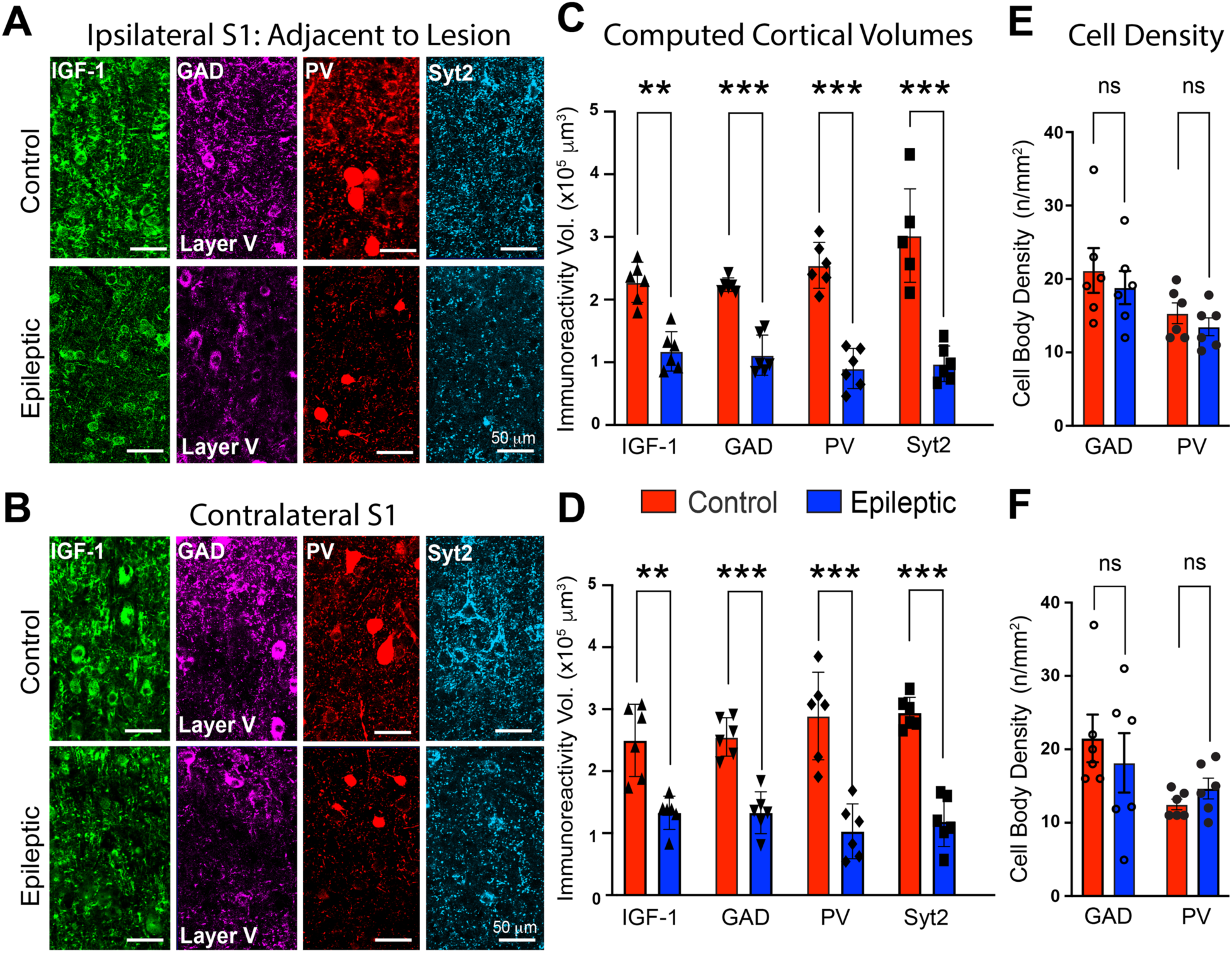
Alterations in the expression of IGF-1, GAD, PV, and synaptotagmin 2. (A and B) High magnification images of area S1 in epileptic rats show a decrease in proteins expressed by putative GABAergic nerve terminals yet the number of immunolabeled interneurons is unchanged. IGF-1 expression is also suppressed in the epileptic neocortex both ipsilateral and contralateral to TTX-induced lesion. Note the relatively sparce putative presynaptic nerve terminal labeling by all three GABAergic proteins in the epileptic neocortex. (C and D) Volumetric quantification of immunoreactivity performed with Imaris surface rendering/reconstruction algorithm. (E and F) Neuron cell body density computed with Neurolucida. N=6 for both groups: 3 males, 3 females. ** p ≤ 0.01, *** p ≤ 0.001, ns non-significant - two-way ANOVA corrected for multiple comparisons. (Reproduced from reference 61)

We next sought to determine whether similar changes take place in children with IESS. To do so, we obtained surgically resected neocortical tissue from IESS patients who had undergone surgery to control their seizures. All had a history of perinatal strokes. For comparison, we examined resected cortical tissue from patients that was adjacent to tumors removed during cancer surgery. Supplemental Figure 2 shows that compared to the tumor control tissue, there was a dramatic decrease in the expression of GAD, PV, and IGF-1 in spasm patients, marked by a decrease in the density of putative GABAergic nerve terminals ^61^.

As mentioned earlier, brain-derived IGF-1 has well-known neuronal growth-promoting properties^57^. Thus, we wondered if our observed reduction in IGF-1 expression in the cortex of epileptic rats and children could impact interneuron development and contribute to the observed decreases in interneuron synaptic proteins and concomitant diminished connectivity. It is well known that GABAergic interneurons undergo dramatic increases in connectivity during the first month of postnatal life in rodents. To address the hypothesis that a reduction of IGF-1 in the epileptic neocortex impairs the development of inhibitory interneurons, we sought to eliminate the receptor for IGF-1 (IGF-1R) from neocortical neurons soon after birth. To accomplish this, we knocked-down IGF-1R expression in conditional IGF-1R^lox^ knockout mice by a unilateral focal injection of AAV-Cre virus into the neonatal neocortex. IGF-1R^lox^ mice injected with control AAV lacking Cre and naïve mice served as controls. Results showed one month later that IGF-1R expression was greatly reduced in neocortical regions expressing the AAV-Cre reporter. Importantly, in these same regions we also observed a reduction in the GAD, PV, and synaptotagmin 2 synaptic networks that had developed by this age ^61^. Thus, a reduction in IGF-1 signaling through its receptor appears to impact the maturation of inhibitory interneurons and GABAergic synaptogenesis. This led us to suspect that the loss of IGF-1 in the epileptic cortex impaired interneuron development, contributing at least in part to the loss of neocortical inhibitory interneuron connectivity.

## (1-3)IGF-1 rescues inhibitory interneuron connectivity, restores IGF-1 expression, and abolishes spasms

To further explore the hypothesis that the loss of IGF-1 brought about diminished synaptic inhibition, we reasoned that experimental strategies that would restore IGF-1 levels might also renew interneuron development, rescue GABAergic synaptic connectivity, and possibly abolish spasms. One way to test this hypothesis would be to treat epileptic animals with IGF-1— essentially, a replacement therapy. While treatment with commercially available recombinant human (rh)IGF-1 was an obvious first choice, results suggested that, due to constraints of the blood-brain barrier, rhIGF-1 was only able to activate the IGF-1 signaling pathway in the neocortex at dosages reported to produce significant peripheral side effects such as hypoglycemia ^61, 73^. Thus, we turned to the IGF-1-derived tripeptide (1-3)IGF-1, which had been reported to have neurotropic properties similar to IGF-1 ^74^ and to cross the blood-brain barrier ^75^. Moreover, our early biochemical studies had provided evidence that (1-3)IGF-1 “acted through” IGF-1R in activating the IGF-1 downstream signaling effectors such as AKT ^47^. However, at that time the exact mechanism was unclear, since it was very unlikely that the tripeptide would bind to the receptor for full length IGF-1 (70 amino acids). Nonetheless, in two reported experimental series, treatment with (1-3)IGF-1 did abolish spasms and hypsarrhythmia in a majority of epileptic animals. In the first, a dosage of 10 mg/kg (i.p.) per day for 21 days abolished spasms in over 60% of animals ^47^. In the second, a higher dosage of 40mg/kg/day for 21 days was given by gavage and abolished spasms in over 80% of animals. In this latter study, daily spasms counts were reduced by over 95% ^61^.

When we compared immunohistochemical results in vehicle- and (1-3)IGF-1-treated rats to those from saline-infused non-epileptic controls, we found that treatment with (1-3)IGF-1 rescued GAD, PV, and synaptotagmin 2 expression to an extent comparable to non-epileptic controls ^61^. Figures 3A and 3B compare images taken from areas S1 to their counterparts in the two control groups and reveal the rescue of the dense network of GABAergic nerve terminals, both ipsilateral and contralateral to the TTX-induced lesion, which is quantified in Figures 3C and 3D, respectively. At this point, our results supported the hypothesis that treatment with (1-3)IGF-1 leads to activation of IGF-1R, which in turn rescues interneuron connectivity and abolishes epileptic spasms. However, it was still unclear how (1-3)IGF-1 activated IGF-1R, but since IGF-1 is the natural agonist for IGF-1R, we examined IGF-1 expression in the epileptic rats treated with the tripeptide and their controls. Results in Figure 4 revealed a restoration of IGF-1 expression, both ipsilateral and contralateral to the TTX lesion. Thus, restoring neocortical levels of IGF-1 by (1-3)IGF-1 is likely responsible for rescuing GABAergic interneuron connectivity and the cessation of spasms.

**Figure 3.**
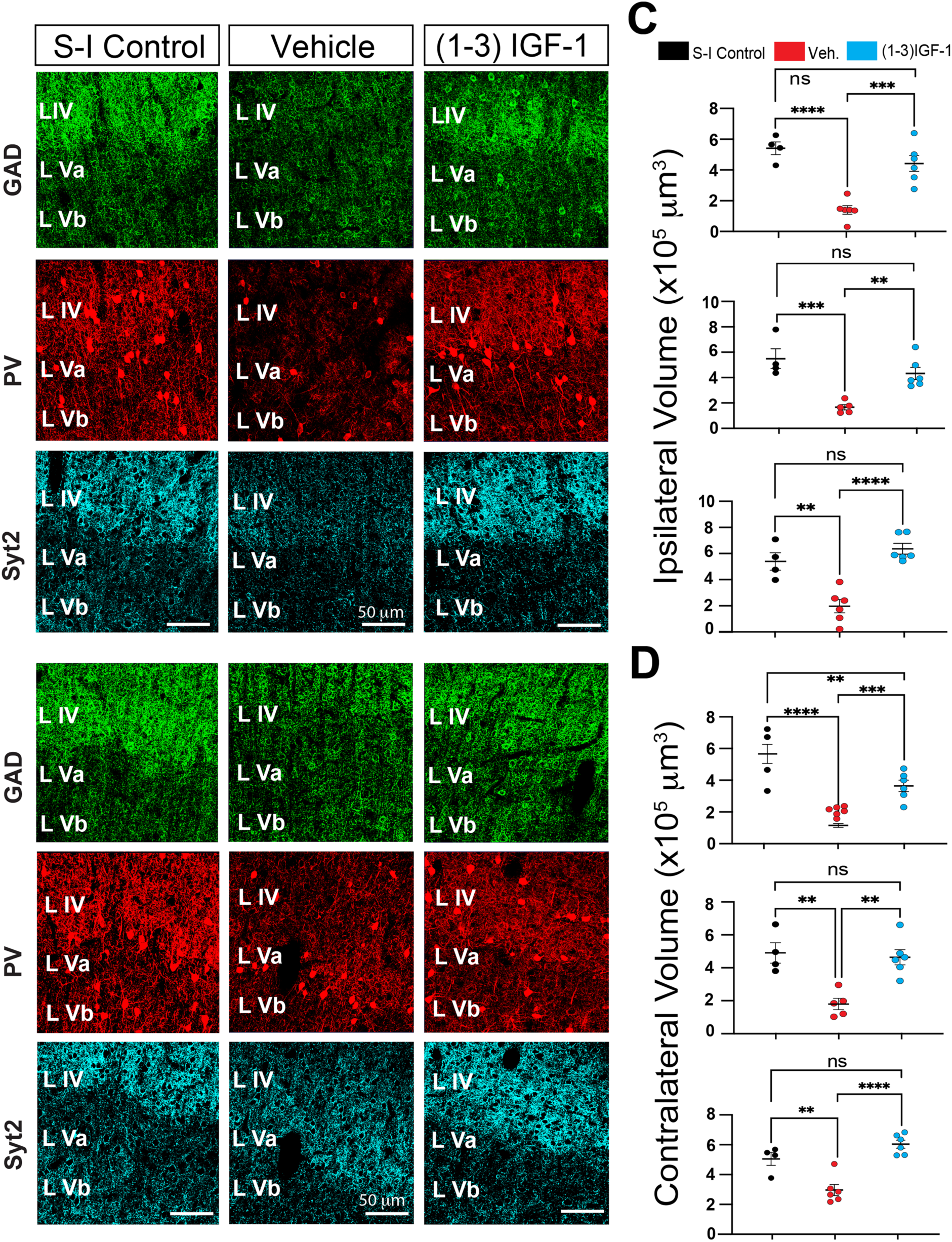
(1-3)IGF-1 rescues neocortical GABAergic nerve terminal networks in epileptic rats. (A and B) Comparison of immunohistochemical results in non-epileptic, saline-infused control rats and epileptic TTX-infused rats treated with either (1-3)IGF-1 or its vehicle. Images from neocortical area S1 (ipsilateral A and contralateral B to infusion site) are shown. Note the density of putative GABAergic nerve terminals is reduced in the epileptic neocortex of animals treated with vehicle, but after (1-3)IGF-1 treatment, presynaptic nerve terminal density approaches that seen in saline-infused controls. (C and D) Volumetric quantification of GAD65/67, PV, and synaptotagmin 2 immunolabeling performed by Imaris surface rendering and reconstruction algorithms. ** p ≤ 0.01, *** p ≤ 0.001, **** p ≤ 0.0001, ns non-significant - one-way ANOVA corrected for multiple comparisons. n = 7 for vehicle, 6 for (1-3)IGF-1 and 4 for controls. S-I – Saline-Infused Control. (Reproduced from reference 61)

**Figure 4.**
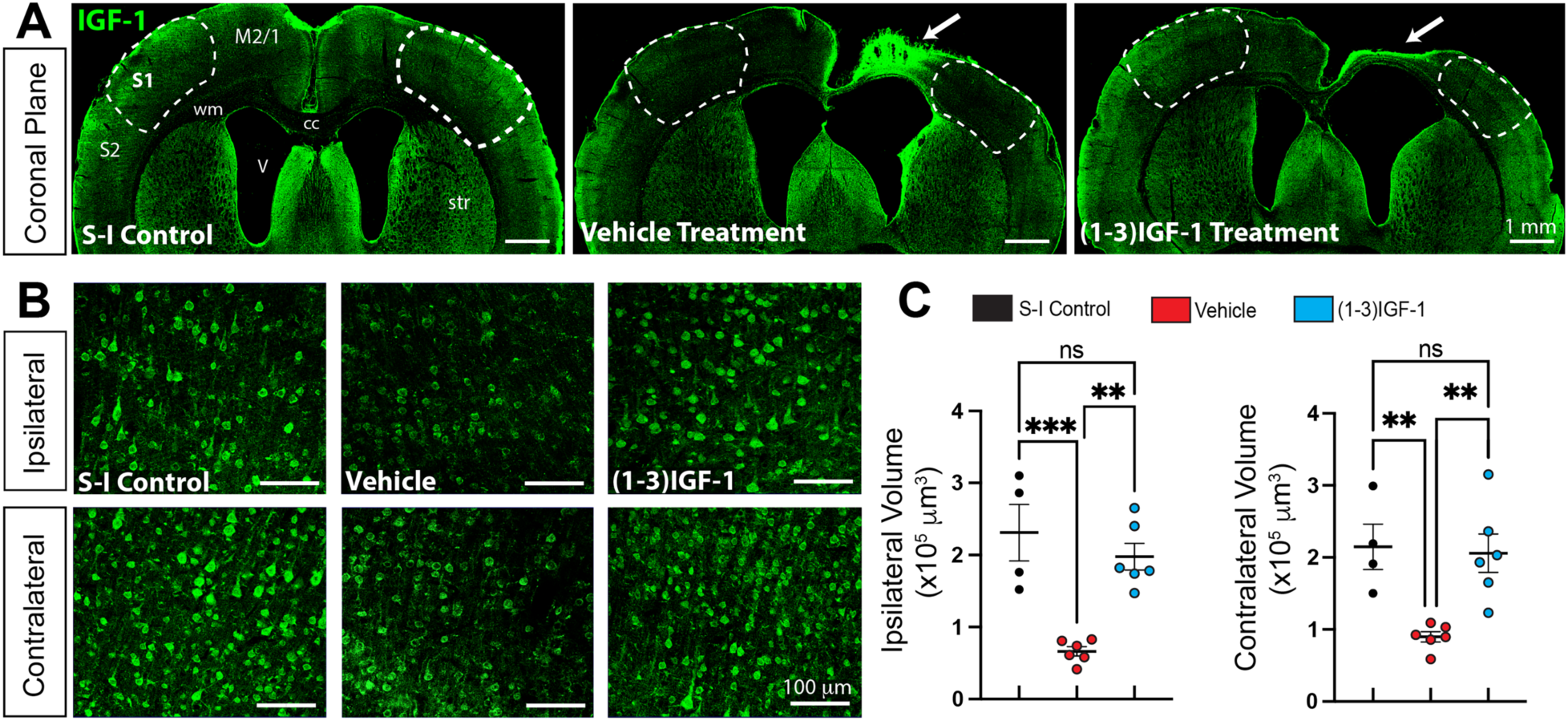
(1-3)IGF-1 restores IGF-1 levels in the neocortex of epileptic rats. (A) IGF-1 immunolabeling in coronal sections from a non-epileptic, saline-infused control rat and TTX-infused epileptic rats treated with either vehicle or (1-3)IGF-1. As reported previously, IGF-1 levels are increased within the TTX lesion (see arrows) but elsewhere in the cortex of vehicle-treated rats, including areas S1 (outlined by dashed lines), IGF-1 expression is suppressed. IGF-1 levels approach that of the saline-infused control after (1-3)IGF-1 treatment. (B) Higher magnification of IGF-1 immunolabeling of area S1. (C) Volumetric quantification of IGF-1 immunolabeling done using Imaris surface rendering and reconstruction algorithms. ** p ≤ 0.01, *** p ≤ 0.001, ns non-significant - one-way ANOVA corrected for multiple comparisons. n = 7 for vehicle, 6 for (1-3)IGF-1 and 4 for controls. Motor Cortex -M2/1, Somatosensory Cortex – S1 and S2, wm – white matter, str – striatum, cc – corpus callosum, V ventricle, S-1 – Saline-Infused Control. (Reproduced from reference 61)

## The inhibitory interneuron dysmaturation hypothesis

When considering possible mechanisms that could explain both injury-induced loss of neocortical interneuron connectivity and its rescue by (1-3)IGF-1, we believe the most likely mechanism is interneuron dysmaturation, or arrested development ^76, 77^. Primary neuronal dysmaturation has been observed in the neocortex ^78, 79^. In these experimental systems, brief periods of prenatal hypoxemia or hypoxia-ischemia resulted in diminished neuronal connectivity, but not neuronal death. But how does brain injury result in dysmaturation? It is well known that brain-derived growth factors, including IGF-1, play important roles in neuron development. Since we observed decreased neocortical expression of IGF-1 in epileptic rats, and the elimination of IGF-1R from neonatal cortices resulted in decreases in interneuron connectivity a few weeks later, the injury-induced loss of IGF-1 is likely an important contributor to inhibitory interneuron dysmaturation. The ability of (1-3)IGF-1 to rescue interneuron connectivity is fully consistent with the dysmaturation hypothesis, especially since the tripeptide restores IGF-1 levels in the cortex which could reinitiate interneuron growth.

Considering the results reviewed above, we have proposed the following schemes for the induction of epileptic spasms and their resolution by (1-3)IGF-1 (Figure 5). For induction, we posit that an injury leads to a reduction in IGF-1 levels in the neocortex which impairs interneuron growth, leading to a reduction in the number of GABAergic synapses. We further hypothesize that diminished synaptic inhibition contributes to the generation of epileptic spasms. As for spasm resolution, we propose that treatment with (1-3)IGF-1 restores IGF-1 levels in the cortex and re-establishes IGF-1R molecular signaling. This reinstates GABAergic synapse formation, restores synaptic inhibition, and abolishes spasms.

**Figure 5.**
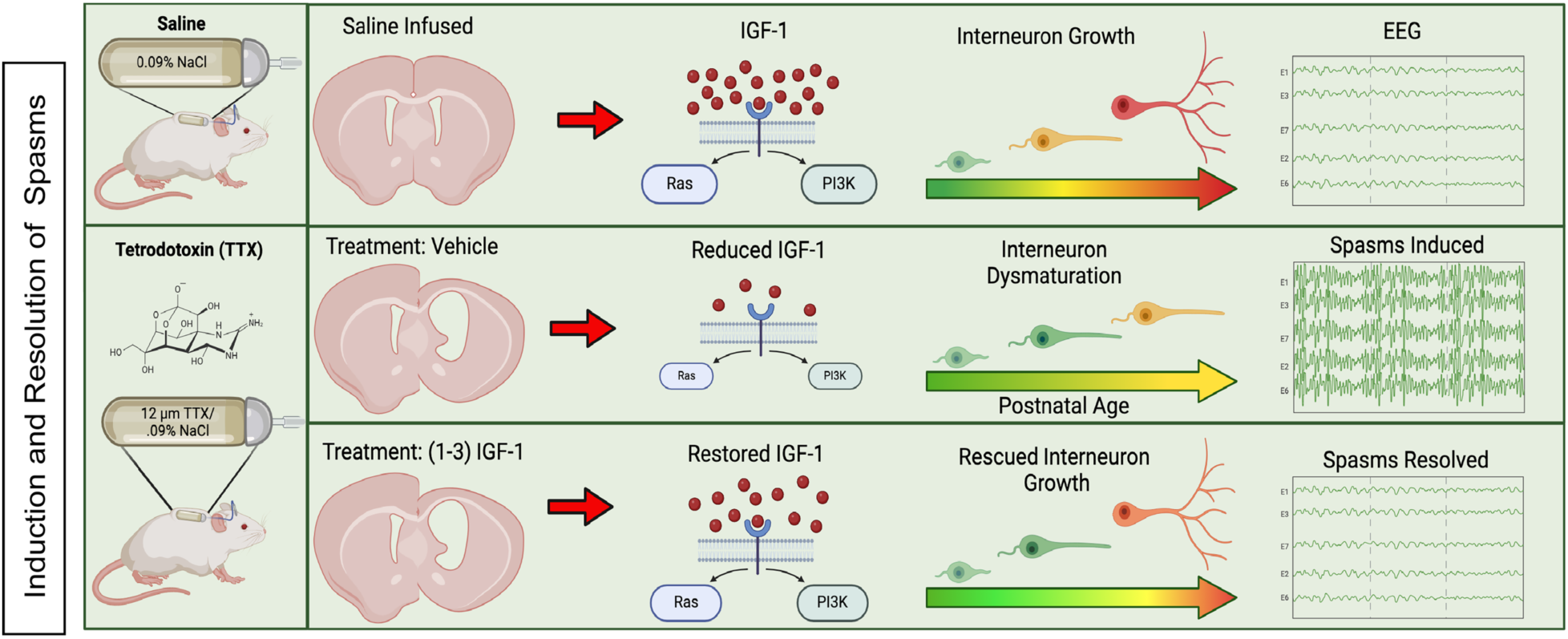
Hypothetical schemes for the role IGF-1 plays in inducing and resolving epileptic spasms. In saline-infused control rats, IGF-1 signaling through the IGF-1R signaling pathways plays a role in interneuron growth. TTX-induced neocortical injury diminishes IGF-1 expression and reduces signaling through its growth pathway, resulting in interneuron dysmaturation (impaired maturation) that likely contributes to the induction of epileptic spasms. Treatment with (1-3)IGF-1 restores IGF-1 levels in the cortex, which activates its growth signaling pathway, rescues interneuron growth, and abolishes epileptic spasms. Created with BioRender.com. (Reproduced from reference 61)

Thus (1-3)IGF-1 or one of its analogues are potentially novel disease-modifying therapies for epileptic spasms. It is important to note that IGF-1 analogues are currently in clinical trials for other developmental brain disorders, and one has been FDA-approved to treat girls with Rett syndrome. That said, these drugs may not be useful in treating every child with spasms. As mentioned earlier, there are over 200 medical conditions associated with infantile spasms ^15^. Riikonen and colleagues reported low CSF levels of IGF-1 only in children with symptomatic spasms ^56^; that is, they had an identified brain abnormality, such as a perinatal stroke, often seen by MRI. The TTX animal model is a brain injury model and while (1-3)IGF-1 is effective in it, it has yet to be shown to be effective for other etiologies and may be etiology-specific.

## Some final thoughts on the often-debated shared final common pathway hypothesis for spasms

As mentioned earlier, both clinicians and research scientists have long tried to understand how so many disparate medical conditions lead to the same unique epileptic syndrome. Many have speculated that there must be a final common pathway shared by all patients, even though the presumed instigating conditions are so radically different. That long-sought shared mechanism could be impaired GABAergic synaptic inhibition. ARX mutations that result in X-linked infantile spasms syndrome are thought to impair the migration of interneurons and reduce the number of inhibitory interneurons and neocortical interneuron connectivity ^68^. Mutations of the GABA receptors that diminish GABA receptor expression and synaptic inhibition (GABRB1, GABRB3 and GABRA1, among others ^80-83^) are all associated with IESS. We propose that neocortical injuries result in neocortical interneuron dysmaturation ^61^. In addition, evidence has accumulated that mutations of TSC1 ^84^ and STXBP1 ^85^, the genes most commonly reported to produce IESS, also result in impaired neocortical synaptic inhibition. This could explain why vigabatrin is effective in treating some patients (e.g., those with TSC mutation). However, since only 25% of IESS patients respond to vigabatrin (and all other drugs that enhance GABAergic inhibition are ineffective), other strategies are needed. These will most likely need to be *etiology specific*—that is, they should not target downstream at the GABAergic synapse itself but the mechanisms upstream of the shared pathway that produced the disinhibition in the first place, such as gene therapy for ARX mutations that supports interneuron migration, gene therapy for STXBP1 to restore GABA release, and (1-3)IGF-1 that targets dysmaturation after neocortical injury. The scheme in Figure 6 summarizes our conceptualization of impaired synaptic inhibition as a shared mechanism for epileptic spasm generation.

**Figure 6.**
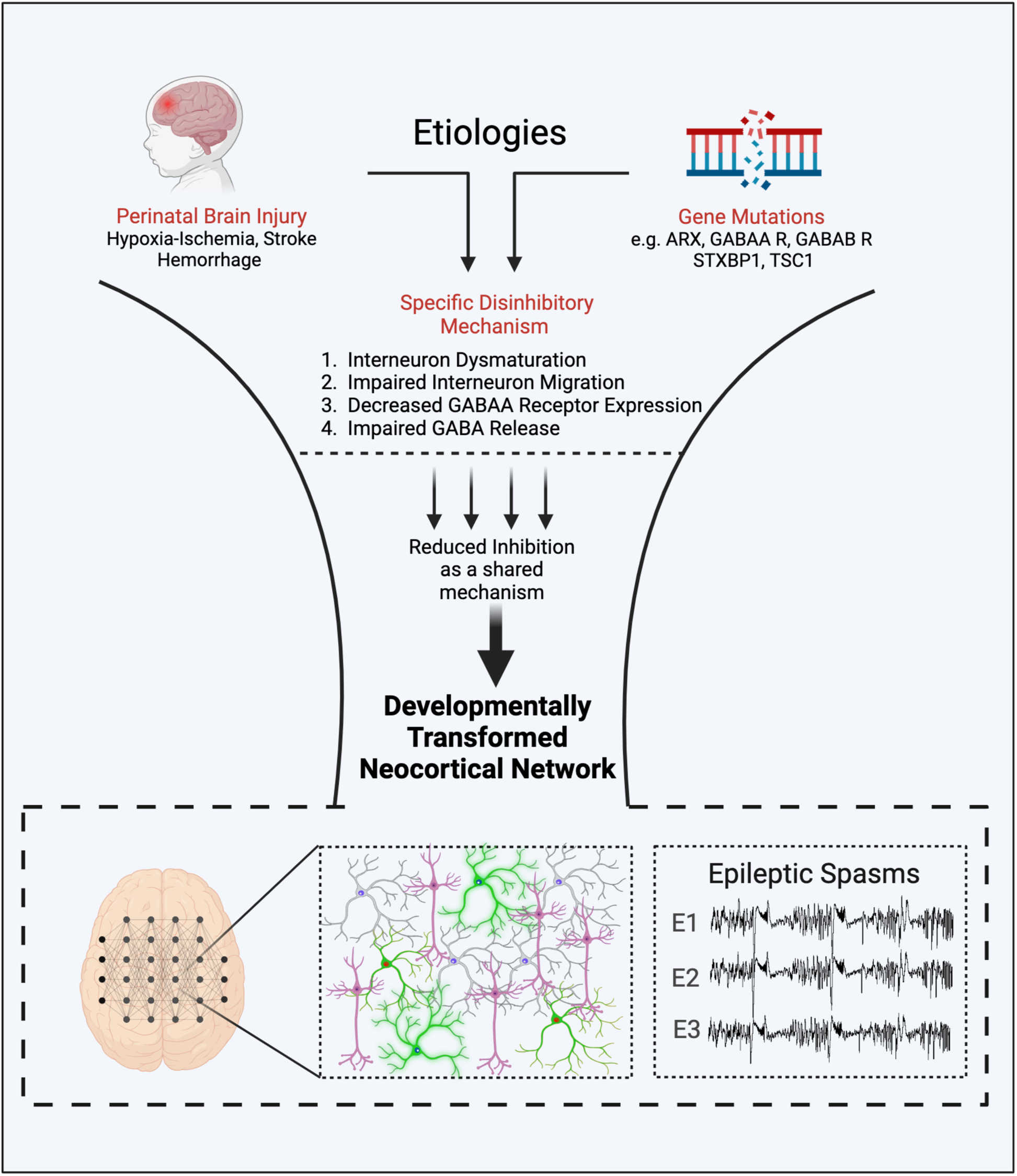
Impaired GABAergic synaptic inhibition as a shared mechanism mediating the generation of epileptic spasms arising from numerous etiologies. While perinatal brain injuries and a variety of gene mutations result in diminished synaptic inhibition, the specific mechanisms underlying the loss of inhibition are very different. For example, while brain injury results in interneuron dysmaturation, mutations of *ARX* prevents interneuron migration into the cortex. Nonetheless, we hypothesize that the impact on neocortical networks would be the same and that reduced synaptic inhibition is a shared mechanism across a variety of etiologies. In addition, we suggest that the neuronal hyperexcitability produced by diminished inhibition gradually transforms neocortical networks in an activity-dependent fashion during a critical period in brain development to produce the unique epileptiform events of epileptic spasms and their behavioral accompaniments. (See ^30^ for demonstration of epileptic spasm progression in the TTX model that might mediate network transformation.) Created with BioRender.com.

Nonetheless, the fact that only one drug designed to target GABAergic inhibition is effective (and only in a minority of IESS patients) could also indicate that mechanisms other than disinhibition produce IESS. Like all forms of epilepsy, epileptic spasms are a neuronal network phenomenon characterized by uncontrolled synchronous discharging of many neurons in a network. It is easy to see how impaired inhibitory synaptic transmission could lead to enhanced network excitability and spasms, but there are many other potential molecular/cellular mechanisms that can produce network hyperexcitability. These might include enhanced glutamatergic synaptic transmission due to mutations of genes for postsynaptic receptors, or molecules involved in presynaptic release of glutamate, or enhanced intrinsic excitability of excitatory neurons due to ion channel mutations, among many others. Thus, there may not be one shared pathway (e.g., disinhibition) all patients have in common but instead multiple shared pathways involved in the generation of spasms (see a proposed scheme in Supplemental Figure 3). However, we propose that each pathway ultimately impinges on a neocortical network and transforms it in an activity-dependent way during a developmental critical period to produce epileptic spasms. *The transformation of a neocortical network to one that produces spasms* (see Figure 1 and Supplemental Figure 1) *would be the only shared mechanism all patients have in common* in the generation of epileptic spasms.

## Acknowledgements

This work was funded by CURE’s Infantile Spasms Initiative and NIH NINDS grants: RO1 NS105913 and R61/R33 NS112553 to J.W.S. It was also supported by IDDRC grant 1U54 HD083092 from the Eunice Kennedy Shriver National Institute of Child Health and Human Development.

## Author Contributions

C.J.B-R., J.W.S., and C.-H. Lee contributed to the conception and design of studies reviewed here, the acquisition and analysis of data, and drafting the text and preparing the figures.

## Figures

**Supplemental Figure 1.**
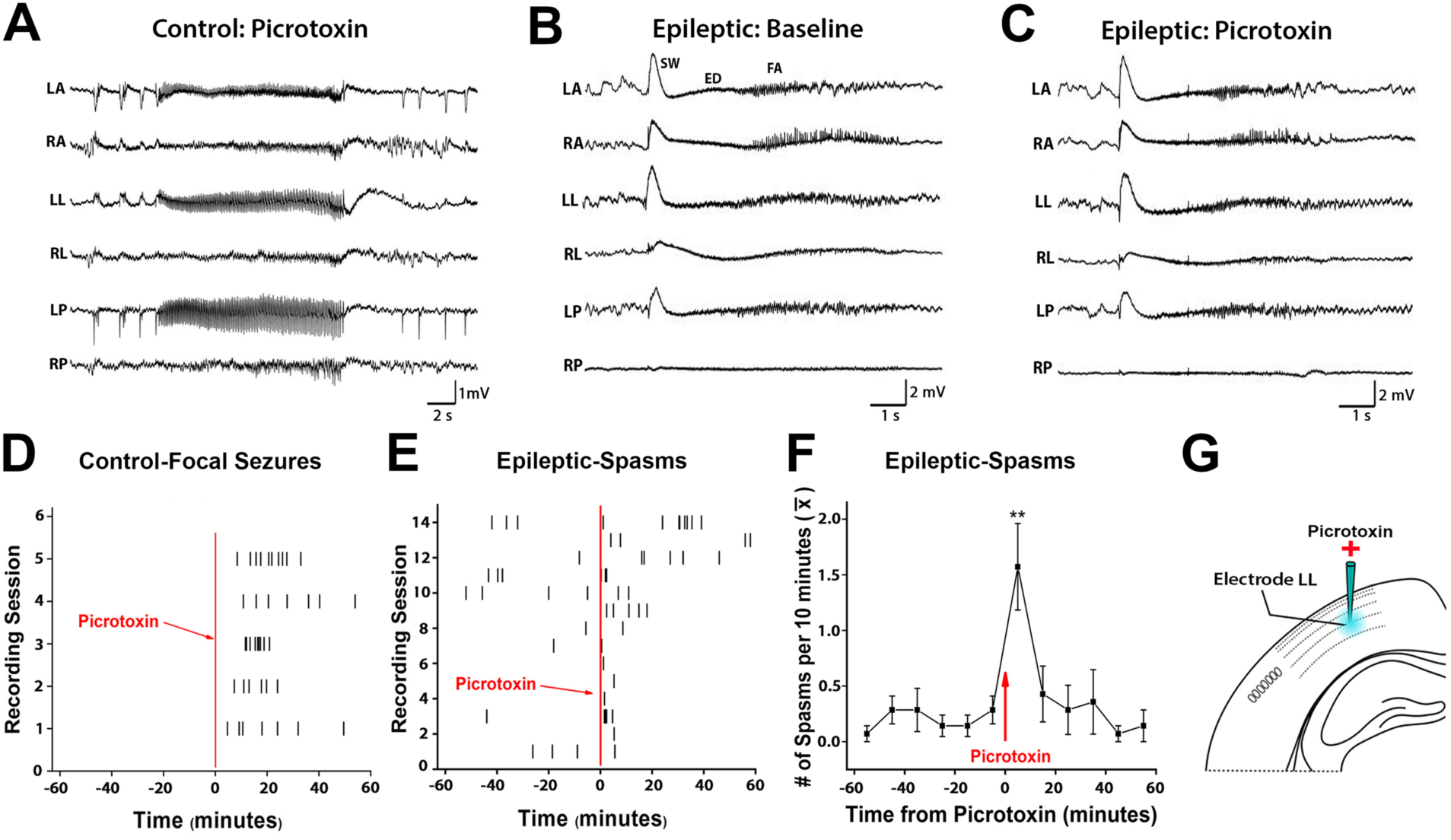
Neocortical infusion of picrotoxin induces focal seizures in control animals but spasms in epileptic animals. (A) Acute picrotoxin infusion into the neocortex of control rats induced recurring focal seizures. (B and C) Spasms were induced in epileptic rats that were identical to those occurring spontaneously during baseline recordings. SW: slow wave of ictal event, ED: electrodecrement, FA: fast activity. (D and E) Raster plots of the occurrence of focal seizures and spasms across 2-hour recordings sessions—1 hour before picrotoxin (i.e., baseline) and 1 hour afterwards. (F) Average spasm frequency across 14 recordings sessions in 10-minute increments (** p ≤ 0.01 compared to all preinfusion times: one-way ANOVA). (G) Schematic illustrating picrotoxin infusion.

**Supplemental Figure 2.**
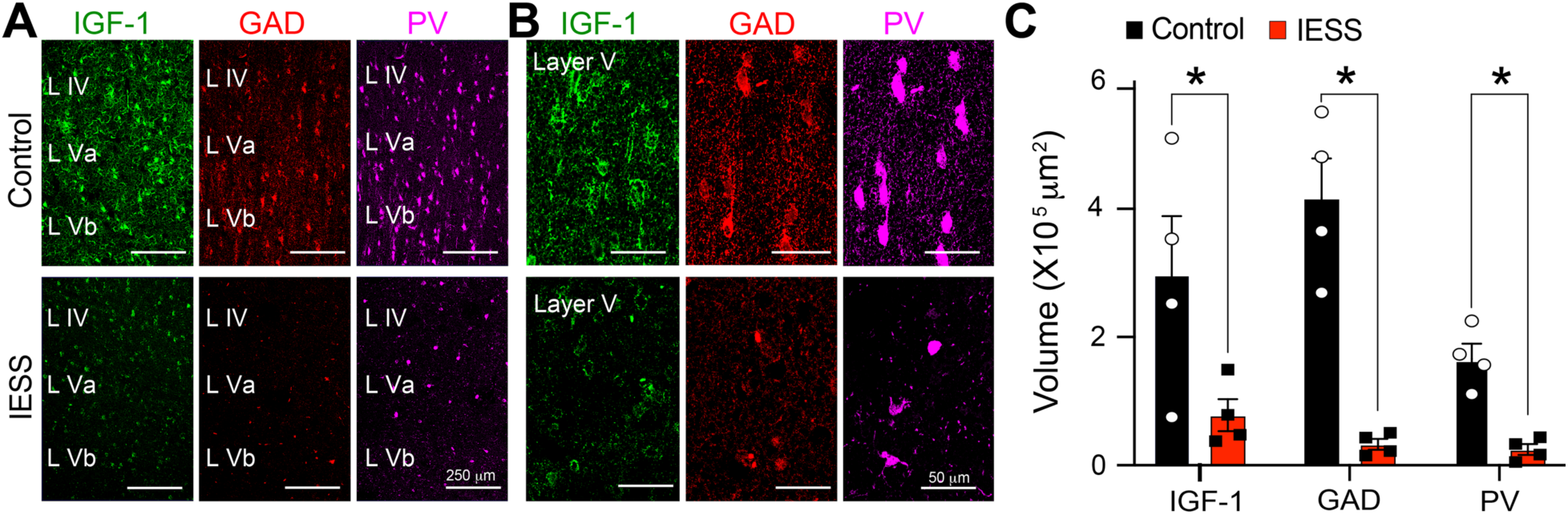
Alterations in the expression of IGF-1, GAD, and PV in the neocortex of IESS patients. (A) Compared to resected neocortical tissue adjacent to tumors in control patients, expression of GAD, PV, and IGF-1 were reduced in surgically resected tissue (adjacent to frank GFAP-positive gliotic neocortical lesions) in IESS patients. Interneuron cell bodies appear noticeably smaller in epileptic tissue. (B) At higher magnification, sparse labeling of putative GABAergic nerve terminals is apparent in sections from spasm patients. (C) Quantification of results with Imaris surface rendering and reconstruction algorithms. * p ≤ 0.05 - two-way ANOVA corrected for multiple comparisons. (Reproduced from reference 61)

**Supplemental Figure 3.**
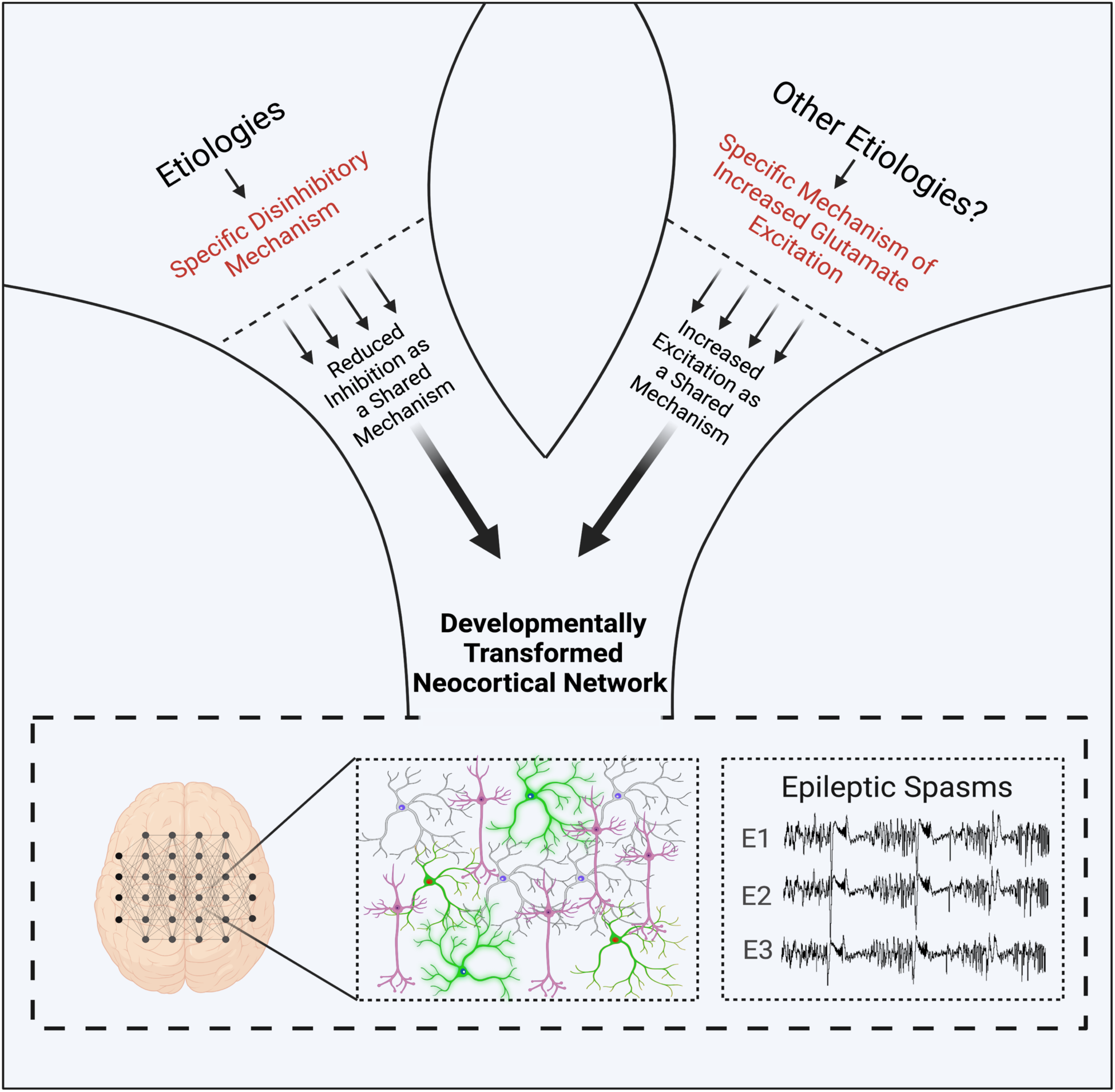
Multiple shared mechanisms may underlie epileptic spasm generation. In addition to diminished inhibition, other specific cellular and molecular mechanisms that increase neocortical network excitability could produce epileptic spasms through other shared mechanisms. In this simplified example, multiple specific mechanisms could increase glutamatergic synaptic transmission in different ways – e.g. increase in glutamatergic synapse number or presynaptic glutamate release etc. However, each mechanism would have in common enhanced excitatory synaptic transmission. Like disinhibition, the resulting neuronal hyperexcitability could gradually transform neocortical networks in an activity-dependent manner during a critical period in development to produce epileptic spasms. Importantly, this model would suggest that the only shared mechanism all patients with spasms have in common is the transformation of neocortical networks into ones with the unique characteristics of spasms. Created with BioRender.com

## References

1. Chevrie JJ, Aicardi J. Convulsive disorders in the first year of life: persistence of epileptic seizures. Epilepsia. 1979 Dec;20(6):643–9.

2. Pellock JM, Hrachovy R, Shinnar S, et al. Infantile spasms: a U.S. consensus report. Epilepsia. 2010 Oct;51(10):2175–89.

3. Zuberi SM, Wirrell E, Yozawitz E, et al. ILAE classification and definition of epilepsy syndromes with onset in neonates and infants: Position statement by the ILAE Task Force on Nosology and Definitions. Epilepsia. 2022 Jun;63(6):1349–97.

4. Jeavons PM, Bower BD. The natural history of infantile spasms. Arch Dis Child. 1961 Feb;36:17–22.

5. Riikonen R. A long-term follow-up study of 214 children with the syndrome of infantile spasms. Neuropediatrics. 1982 Feb;13(1):14–23.

6. Jeavons PM, Bower BD, Dimitrakoudi M. Long-term prognosis of 150 cases of "West syndrome". Epilepsia. 1973 Jun;14(2):153–64.

7. Kellaway P, Hrachovy RA, Frost JD, Jr., Zion T. Precise characterization and quantification of infantile spasms. AnnNeurol. 1979 9/1979;6(3):214-8.

8. Gibbs EL, Fleming MM, Gibbs FA. Diagnosis and prognosis of hypsarhythmia and infantile spasms. Pediatrics. 1954 1/1954;13(1):66-73.

9. Hrachovy RA, Frost JD, Jr., Kellaway P. Hypsarrhythmia: variations on the theme. Epilepsia. 1984 6/1984;25(3):317-25.

10. Mackay MT, Weiss SK, ms-Webber T, et al. Practice parameter: medical treatment of infantile spasms: report of the American Academy of Neurology and the Child Neurology Society. Neurology. 2004 5/25/2004;62(10):1668-81.

11. Hussain SA, Shinnar S, Kwong G, et al. Treatment of infantile spasms with very high dose prednisolone before high dose adrenocorticotropic hormone. Epilepsia. 2014 Jan;55(1):103–7.

12. Willmore LJ, Abelson MB, Ben-Menachem E, Pellock JM, Shields WD. Vigabatrin: 2008 update. Epilepsia. 2009 2/2009;50(2):163-73.

13. Knupp KG, Coryell J, Nickels KC, et al. Response to treatment in a prospective national infantile spasms cohort. Ann Neurol. 2016 Mar;79(3):475–84.

14. Shields WD. Infantile Spasms: Little Seizures, BIG Consequences. Epilepsy Curr. 2006 5/2006;6(3):63-9.

15. Frost Jr. JD, Hrachovy RA. Infantile Spasms. Boston: Kluber Academic Publishers; 2003.

16. Wirrell EC, Shellhaas RA, Joshi C, et al. How should children with West syndrome be efficiently and accurately investigated? Results from the National Infantile Spasms Consortium. Epilepsia. 2015 Apr;56(4):617–25.

17. Scantlebury MH, Galanopoulou AS, Chudomelova L, Raffo E, Betancourth D, Moshe SL. A model of symptomatic infantile spasms syndrome. Neurobiol Dis. 2010 Mar;37(3):604–12.

18. Marsh E, Fulp C, Gomez E, et al. Targeted loss of Arx results in a developmental epilepsy mouse model and recapitulates the human phenotype in heterozygous females. Brain. 2009 6/2009;132(Pt 6):1563-76.

19. Cortez MA, Shen L, Wu Y, et al. Infantile Spasms and Down syndrome: A New Animal Model. PediatrRes. 2009 1/28/2009.

20. Price MG, Yoo JW, Burgess DL, et al. A triplet repeat expansion genetic mouse model of infantile spasms syndrome, Arx(GCG)10+7, with interneuronopathy, spasms in infancy, persistent seizures, and adult cognitive and behavioral impairment. JNeurosci. 2009 7/8/2009;29(27):8752-63.

21. Qu S, Jackson LG, Zhou C, et al. Heterozygous GABA(A) receptor beta3 subunit N110D knock-in mice have epileptic spasms. Epilepsia. 2023 Apr;64(4):1061–73.

22. Velisek L, Jehle K, Asche S, Veliskova J. Model of infantile spasms induced by N-methyl-D-aspartic acid in prenatally impaired brain. AnnNeurol. 2007 2/2007;61(2):109-19.

23. Pirone A, Alexander J, Lau LA, et al. APC conditional knock-out mouse is a model of infantile spasms with elevated neuronal beta-catenin levels, neonatal spasms, and chronic seizures. Neurobiol Dis. 2017 Feb;98:149–57.

24. Lee CL, Frost JD, Jr., Swann JW, Hrachovy RA. A new animal model of infantile spasms with unprovoked persistent seizures. Epilepsia. 2008 2/2008;49(2):298-307.

25. Galvan CD, Hrachovy RA, Smith KL, Swann JW. Blockade of neuronal activity during hippocampal development produces a chronic focal epilepsy in the rat. JNeurosci. 2000 2000;20(8):2904–16.

26. Galvan CD, Wenzel JH, Dineley KT, et al. Postsynaptic contributions to hippocampal network hyperexcitability induced by chronic activity blockade in vivo. EurJNeurosci. 2003 10/2003;18(7):1861-72.

27. Frost JD, Jr., Lee CL, Le JT, Hrachovy RA, Swann JW. Interictal high frequency oscillations in an animal model of infantile spasms. NeurobiolDis. 2012 5/2012;46(2):377-88.

28. Le JT, Frost JD, Jr., Swann JW. Acthar(R) Gel (repository corticotropin injection) dose-response relationships in an animal model of epileptic spasms. Epilepsy & behavior: E&B. 2021 Mar;116:107786.

29. Frost JD, Jr., Le JT, Lee CL, Ballester-Rosado C, Hrachovy RA, Swann JW. Vigabatrin therapy implicates neocortical high frequency oscillations in an animal model of infantile spasms. Neurobiol Dis. 2015 Oct;82:1–11.

30. Le JT, Ballester-Rosado CJ, Frost JD, Jr., Swann JW. Neurobehavioral deficits and a progressive ictogenesis in the tetrodotoxin model of epileptic spasms. Epilepsia. 2022 Dec;63(12):3078–89.

31. Semple BD, Blomgren K, Gimlin K, Ferriero DM, Noble-Haeusslein LJ. Brain development in rodents and humans: Identifying benchmarks of maturation and vulnerability to injury across species. Prog Neurobiol. 2013 Jul-Aug;106-107:1-16.

32. Stafstrom CE. Infantile spasms: a critical review of emerging animal models. Epilepsy Curr. 2009 5/2009;9(3):75-81.

33. de Menezes MA, Rho JM. Clinical and electrographic features of epileptic spasms persisting beyond the second year of life. Epilepsia. 2002 Jun;43(6):623–30.

34. Camfield P, Camfield C, Lortie A, Darwish H. Infantile spasms in remission may reemerge as intractable epileptic spasms. Epilepsia. 2003 Dec;44(12):1592–5.

35. Talwar D, Baldwin MA, Hutzler R, Griesemer DA. Epileptic spasms in older children: persistence beyond infancy. Epilepsia. 1995 Feb;36(2):151–5.

36. Chugani HT, Shields WD, Shewmon DA, Olson DM, Phelps ME, Peacock WJ. Infantile spasms: I. PET identifies focal cortical dysgenesis in cryptogenic cases for surgical treatment. AnnNeurol. 1990 4/1990;27(4):406-13.

37. Chipaux M, Dorfmuller G, Fohlen M, et al. Refractory spasms of focal onset-A potentially curable disease that should lead to rapid surgical evaluation. Seizure. 2017 Oct;51:163–70.

38. Gettings JV, Shafi S, Boyd J, et al. The Epilepsy Surgery Experience in Children With Infantile Epileptic Spasms Syndrome at a Tertiary Care Center in Canada. J Child Neurol. 2023 Mar;38(3-4):113–20.

39. Kolosky T, Goldstein Shipper A, Sun K, et al. Epilepsy surgery for children with epileptic spasms: A systematic review and meta-analysis with focus on predictors and outcomes. Epilepsia Open. 2024 Aug;9(4):1136–47.

40. Chugani HT, Shewmon DA, Shields WD, et al. Surgery for intractable infantile spasms: neuroimaging perspectives. Epilepsia. 1993 Jul-Aug;34(4):764-71.

41. Iwatani Y, Kagitani-Shimono K, Tominaga K, et al. Long-term developmental outcome in patients with West syndrome after epilepsy surgery. Brain Dev. 2012 Oct;34(9):731–8.

42. Kang JW, Rhie SK, Yu R, et al. Seizure outcome of infantile spasms with focal cortical dysplasia. Brain Dev. 2013 Sep;35(8):816–20.

43. Chugani HT, Ilyas M, Kumar A, et al. Surgical treatment for refractory epileptic spasms: The Detroit series. Epilepsia. 2015 Dec;56(12):1941–9.

44. Chugani HT, Shewmon DA, Sankar R, Chen BC, Phelps ME. Infantile spasms: II. Lenticular nuclei and brain stem activation on positron emission tomography. AnnNeurol. 1992 2/1992;31(2):212-9.

45. Cohen AL, Mulder BPF, Prohl AK, et al. Tuber Locations Associated with Infantile Spasms Map to a Common Brain Network. Ann Neurol. 2021 Apr;89(4):726–39.

46. Chugani HT. Pathophysiology of infantile spasms. Adv Exp Med Biol. 2002;497:111–21.

47. Ballester-Rosado CJ, Le JT, Lam TT, et al. A Role for Insulin-like Growth Factor 1 in the Generation of Epileptic Spasms. Ann Neurol. 2022 Apr 25.

48. Heck N, Golbs A, Riedemann T, Sun JJ, Lessmann V, Luhmann HJ. Activity-dependent regulation of neuronal apoptosis in neonatal mouse cerebral cortex. Cereb Cortex. 2008 Jun;18(6):1335–49.

49. Leveille F, Papadia S, Fricker M, et al. Suppression of the intrinsic apoptosis pathway by synaptic activity. J Neurosci. 2010 Feb 17;30(7):2623–35.

50. Lee CH, Le JT, Ballester-Rosado CJ, Anderson AE, Swann JW. Neocortical Slow Oscillations Implicated in the Generation of Epileptic Spasms. Ann Neurol. 2021 Feb;89(2):226–41.

51. Alexander GM, Rogan SC, Abbas AI, et al. Remote control of neuronal activity in transgenic mice expressing evolved G protein-coupled receptors. Neuron. 2009 Jul 16;63(1):27–39.

52. Nariai H, Matsuzaki N, Juhasz C, et al. Ictal high-frequency oscillations at 80-200 Hz coupled with delta phase in epileptic spasms. Epilepsia. 2011 10/2011;52(10):e130-e4.

53. Hrachovy RA, Frost JD, Jr., Kellaway P. Sleep characteristics in infantile spasms. Neurology. 1981 6/1981;31(6):688-93.

54. Avanzini G, Panzica F, Franceschetti S. Brain maturational aspects relevant to pathophysiology of infantile spasms. Int Rev Neurobiol. 2002;49:353–65.

55. Lado FA, Moshe SL. Role of subcortical structures in the pathogenesis of infantile spasms: what are possible subcortical mediators? IntRevNeurobiol. 2002 2002;49:115–40.

56. Riikonen RS, Jaaskelainen J, Turpeinen U. Insulin-like growth factor-1 is associated with cognitive outcome in infantile spasms. Epilepsia. 2010 7/2010;51(7):1283-9.

57. O’Kusky J, Ye P. Neurodevelopmental effects of insulin-like growth factor signaling. Front Neuroendocrinol. 2012 Aug;33(3):230–51.

58. Popken GJ, Hodge RD, Ye P, et al. In vivo effects of insulin-like growth factor-I (IGF-I) on prenatal and early postnatal development of the central nervous system. Eur J Neurosci. 2004 Apr;19(8):2056–68.

59. Liu W, Ye P, O’Kusky JR, D’Ercole AJ. Type 1 insulin-like growth factor receptor signaling is essential for the development of the hippocampal formation and dentate gyrus. Journal of neuroscience research. 2009 Oct;87(13):2821-32.

60. Bondy C, Lee WH. Correlation between insulin-like growth factor (IGF)-binding protein 5 and IGF-I gene expression during brain development. J Neurosci. 1993 Dec;13(12):5092–104.

61. Ballester-Rosado CJ, Le JT, Lam TT, Anderson AE, Frost JD, Jr., Swann JW. IGF-1 impacts neocortical interneuron connectivity in epileptic spasm generation and resolution. Neurotherapeutics. 2024 Nov 8;22(1):e00477.

62. Cao P, Maximov A, Sudhof TC. Activity-dependent IGF-1 exocytosis is controlled by the Ca(2+)-sensor synaptotagmin-10. Cell. 2011 Apr 15;145(2):300–11.

63. Garcia-Estrada J, Garcia-Segura LM, Torres-Aleman I. Expression of insulin-like growth factor I by astrocytes in response to injury. Brain Res. 1992 Oct 2;592(1-2):343–7.

64. Madathil SK, Evans HN, Saatman KE. Temporal and regional changes in IGF-1/IGF-1R signaling in the mouse brain after traumatic brain injury. J Neurotrauma. 2010 Jan;27(1):95–107.

65. Ito M, Mikawa H, Taniguchi T. Cerebrospinal fluid GABA levels in children with infantile spasms. Neurology. 1984 Feb;34(2):235–8.

66. Loscher W, Siemes H. Cerebrospinal fluid gamma-aminobutyric acid levels in children with different types of epilepsy: effect of anticonvulsant treatment. Epilepsia. 1985 Jul-Aug;26(4):314-9.

67. Airaksinen E, Tuomisto L, Riikonen R. The concentrations of GABA, 5-HIAA and HVA in the cerebrospinal fluid of children with infantile spasms and the effects of ACTH treatment. Brain Dev. 1992 Nov;14(6):386-90.

68. Kitamura K, Yanazawa M, Sugiyama N, et al. Mutation of ARX causes abnormal development of forebrain and testes in mice and X-linked lissencephaly with abnormal genitalia in humans. NatGenet. 2002 11/2002;32(3):359-69.

69. Kato M, Dobyns WB. X-linked lissencephaly with abnormal genitalia as a tangential migration disorder causing intractable epilepsy: proposal for a new term, "interneuronopathy". JChild Neurol. 2005 4/2005;20(4):392-7.

70. Ryner RF, Derera ID, Armbruster M, et al. Cortical Parvalbumin-Positive Interneuron Development and Function Are Altered in the APC Conditional Knockout Mouse Model of Infantile and Epileptic Spasms Syndrome. J Neurosci. 2023 Feb 22;43(8):1422–40.

71. Katsarou AM, Li Q, Liu W, Moshe SL, Galanopoulou AS. Acquired parvalbumin-selective interneuronopathy in the multiple-hit model of infantile spasms: A putative basis for the partial responsiveness to vigabatrin analogs? Epilepsia Open. 2018 Dec;3(Suppl Suppl 2):155-64.

72. Cortez MA, Shen L, Wu Y, et al. Infantile spasms and Down syndrome: a new animal model. PediatrRes. 2009 5/2009;65(5):499-503.

73. Richmond EJ, Rogol AD. Recombinant human insulin-like growth factor-I therapy for children with growth disorders. Adv Ther. 2008 Dec;25(12):1276–87.

74. Tropea D, Kreiman G, Lyckman A, et al. Gene expression changes and molecular pathways mediating activity-dependent plasticity in visual cortex. NatNeurosci. 2006 5/2006;9(5):660-8.

75. Baker AM, Batchelor DC, Thomas GB, et al. Central penetration and stability of N-terminal tripeptide of insulin-like growth factor-I, glycine-proline-glutamate in adult rat. Neuropeptides. 2005 Apr;39(2):81–7.

76. Volpe JJ. Primary neuronal dysmaturation in preterm brain: Important and likely modifiable. J Neonatal Perinatal Med. 2021;14(1):1–6.

77. Dean JM, Bennet L, Back SA, McClendon E, Riddle A, Gunn AJ. What brakes the preterm brain? An arresting story. Pediatr Res. 2014 Jan;75(1-2):227–33.

78. Dean JM, McClendon E, Hansen K, et al. Prenatal cerebral ischemia disrupts MRI-defined cortical microstructure through disturbances in neuronal arborization. Science translational medicine. 2013 Jan 16;5(168):168ra7.

79. McClendon E, Shaver DC, Degener-O’Brien K, et al. Transient Hypoxemia Chronically Disrupts Maturation of Preterm Fetal Ovine Subplate Neuron Arborization and Activity. J Neurosci. 2017 Dec 6;37(49):11912–29.

80. Monfrini E, Borellini L, Zirone E, et al. GABRB1-related early onset developmental and epileptic encephalopathy: Clinical trajectory and novel de novo mutation. Epileptic Disord. 2023 Dec;25(6):867–73.

81. Farnaes L, Nahas SA, Chowdhury S, et al. Rapid whole-genome sequencing identifies a novel GABRA1 variant associated with West syndrome. Cold Spring Harb Mol Case Stud. 2017 Sep;3(5).

82. Kodera H, Ohba C, Kato M, et al. De novo GABRA1 mutations in Ohtahara and West syndromes. Epilepsia. 2016 Apr;57(4):566–73.

83. Janve VS, Hernandez CC, Verdier KM, Hu N, Macdonald RL. Epileptic encephalopathy de novo GABRB mutations impair gamma-aminobutyric acid type A receptor function. Ann Neurol. 2016 May;79(5):806–25.

84. Fu C, Cawthon B, Clinkscales W, Bruce A, Winzenburger P, Ess KC. GABAergic interneuron development and function is modulated by the Tsc1 gene. Cereb Cortex. 2012 Sep;22(9):2111–9.

85. Chen W, Cai ZL, Chao ES, et al. Stxbp1/Munc18-1 haploinsufficiency impairs inhibition and mediates key neurological features of STXBP1 encephalopathy. Elife. 2020 Feb 19;9.

